# Sophocarpine inhibits TRP channels to produce anti-pruritic and analgesic effects in a mouse model of inflammatory itch and pain

**DOI:** 10.1101/2023.10.12.561966

**Authors:** Hekun Zeng, Peiyang Li, Dan Zhou, Zhe Zhang, Alexei Verkhratsky, Hong Nie

## Abstract

**Background and objective:** Itch, an unpleasant sensation prompting the urge to scratch, and pain, aimed at detecting potential harm through acute withdrawal or protective behaviors, are increasingly recognized as interconnected phenomena. The co-occurrence of itch and pain symptoms in various diseases impairs therapeutic efficacy and the quality of life. In this study, we investigated the potential antipruritic and analgesic effects of sophocarpine (SC), an active compound of *Sophorae Flavesentis Radix*, in a murine model of inflammatory itch and pain, and sought to elucidate the underlying mechanisms.

**Method:** The anti-pruritic and analgesic effects of three doses of SC (60 mg/kg, 30 mg/kg, 10 mg/kg) were tested by analyzing the scratching and wiping behaviors in squaric acid dibutylester (SABDE)-induced allergic contact dermatitis (ACD) mouse model accompany by itch and pain, respectively. Psoriasis area and severity index (PASI) score was used to test the anti-inflammatory effect of SC. The underlying mechanisms were studied by real-time quantitative polymerase chain reaction (RT-qPCR) and western blotting. Additionally, the anti-pruritic and analgesic effects of SC were further tested in mice with intradermal injection of allyl-isothiocyanate (AITC), a TRPA1 agonist, or capsaicin (CAP), a TRPV1 agonist, respectively. The relationships between SC, AITC, CAP and TRPV1, TRPA1 were simulated by molecular docking.

**Results:** SC treatment significantly decreased scratching bouts and wipes, as well as the PASI score. Administration of SC reduced the mRNA and protein expression of both TRPA1 and TRPV1. Moreover, pretreatment of SC decreased scratching bouts and wipes induced by AITC as well as by CAP. Molecular docking revealed potential competitive binding between SC and AITC on TRPA1, and SC and CAP on TRPV1.

**Conclusion:** We demonstrated that SC has strong anti-pruritic and analgesic effects by targeting the TRPA1 and TRPV1 ion channels, and is a potential competitive inhibitor of TRPA1 and TRPV1. These findings suggest that SC has significant therapeutic potential in the therapy of diseases with inflammatory itch and pain.

## 1. Introduction

Itch is an unpleasant sensation that causes the need to scratch the affected area to remove puritogens thus causing temporary relief (1). Differing from the itch, pain-related harm detection instigates acute withdrawal or other protective behaviors. Itch and pain are distinct sensations both relying on interacting sensory subdivisions; however, opioids could inhibit pain but generate itch indicating distinct mechanisms for itch and pain (2).

Chronic itch and pain have broadly overlapping mediators and functions (3). For example, nerve growth factor could induce sensitization of primary afferents in both chronic itch and pain (4). Finding effective medicine targeting the common mechanism to relieve itch and pain is required to avoid physical or mental issues brought by both uncomfortable sensations coexisting.

In humans, the transient receptor potential (TRP) channel family covers 27 members classified into six groups of TRPC (canonical), TRPV (vanilloid), TRPM (melastatin), TRPP (polycystin), TRPML (mucolipin), and TRPA (ankyrin); an additional group of TRPN (NOMPC-like) is present in invertebrates and fishes. The TRP channels are widely expressed through the nervous system and contribute in particular to all types of sensing including thermal sensation, nociception, chemisorption, equilibrioception and taste (5). The TRP channels are archetypal cationic channels, many of which are highly permeable to Ca^2+^ ions (6). The TRPA1 and TRPV1 subfamilies contribute to both inflammatory itch and pain sensation (7). Many pro-inflammatory mediators such as bradykinin, leukotrienes, tumour necrosis factor-α (TNF-α), and interleukin-1β (IL-1β) can activate the TRPA1 and TRPV1 channels to mediate pain (8). In addition, TRPA1 was suggested to mediate histamine-independent itch, whereas TRPV1 mediate histamine-dependent itch (9, 10).

*Sophorae Flavesentis Radix* (SFR), a Traditional Chinese Medicine, has been used to relieve itch and pain in clinics in China for thousands of years. In our previous study, we identified, by cell membrane chromatography, sophocarpine (SC) as a bioactive compound that could bind to receptors on the plasma membrane of cells in dorsal root ganglion (DRG) (11). In the present study, we employed an allergic contact dermatitis (ACD) mouse model of itch and pain to investigate the analgesic and anti-pruritic effect of SC and underlying pathways, and test its effect on the TRPA1 and TRPV1, mediating itch and pain.

## 2. Materials and Methods

### 2.1 Animal

Male mice (6-week-old, 20 ± 2 g) were purchased from the SPF (Beijing) Biotechnology Co. Ltd. (Certificate NO. SCXK (Beijing) 2016-0002). All mice were housed under a cycle of 12 h light/dark and given standard laboratory food and tap water in the Jinan University Medical School’s Laboratory Animal Management Center (Certificate No. SYXK (Guangdong) 2017-0174). All experimental procedures and animal welfare strictly followed the Guide for the Care and Use of Laboratory Animals and related to Jinan University’s ethical regulations (Approval No: IACUC-20180621-07).

### 2.2 Establishment of allergic contact dermatitis mouse model

The ACD, or Contact Hypersensitivity (CHS) mouse model was established as described by applying squaric acid dibutylester (SADBE, Tokyo Chemical Inc., Tokyo, Japan) dissolved in acetone (Guangzhou Chemical Reagent Factory, Guangzhou, China) on the right cheek (12) (Figure 1). Briefly, the mice’s abdomen (1.5×1.5 cm^2^) was shaved on Day -1. Starting from Day 1, 25 µL 1% SADBE (v/v) was applied to the abdominal shaved area for three consecutive days once daily. The mice’s right cheek (0.8×0.8 cm^2^) was shaved on Day 6, and 25 µL 1% SADBE was applied to the cheek shaved area once daily for three consecutive days starting from Day 8 to excite the ACD. The control group was treated with 25 µL pure acetone instead of 1% SADBE.

**Figure 1.**
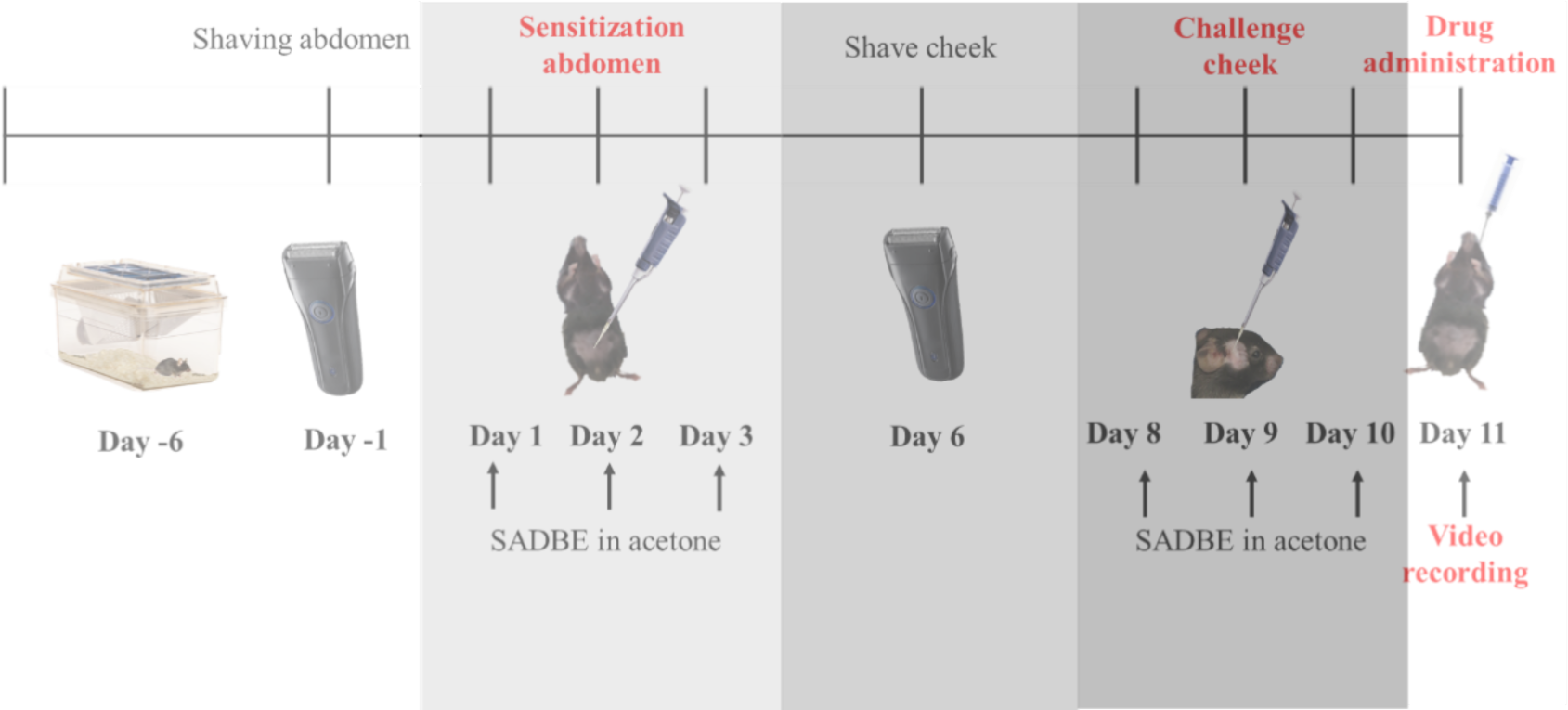
Experimental protocol for the establishment of ACD model on the cheek and drug intragastric administration. 70 C57BL/6 mice were randomly allocated into 7 groups: control group, ACD group, DEM group (5 mg/kg), CPZ group (30 mg/kg), and three doses of SC groups (60 mg/kg, 30 mg/kg, 10 mg/kg). Mice were trained daily from Day 1 to Day 10. On Day -1, the abdomen of the mice was shaved. From Day 1 to Day 3, 25 µL 1% SABDE was applied once daily onto the abdomens, except for the control group of mice who were applied with the same volume of acetone instead. On Day 6, the cheek of the mice was shaved. From Day 8 to Day 10, 25 µL 1% SABDE was applied onto the cheek once daily, except for the control group of mice who were applied with the same volume of acetone instead. On Day 11, mice in the control group and the ACD group were administrated i.g. with saline, the CPZ group with 30 mg/kg CPZ, and the SC groups with corresponding doses of SC. The behaviors were recorded for 2 h.

### 2.3 Drug administration and evaluation of spontaneous scratching and wiping behaviors

Three doses of SC (60, 30 and 10 mg/kg, Chengdu Push Bio-technology Co., Ltd, Chengdu, China), capsazepine (CPZ, 30 mg/kg, Selleck Chemicals LLC, Houston, TX, USA), dexamethasone (DEM, 5 mg/kg, Shanghai Macklin Biochemical Co., Ltd., Shanghai, China) and saline were administrated intragastrically (i.g.) through the catheter on Day 11 (24 h after the third application of 1% SADBE on the cheek) as shown in figure 1. To evaluate the spontaneous scratching bout and wiping behavior, mice were placed in a separate, clear plastic container in 9×9×13 cm^3^ with a small amount of bedding inside, and a camcorder was positioned above the container to record two mice at a time for 2 h, at 1 h after drug administration. Four mirrors were placed with their bottoms aligned to the 4 lines of the square bottom of each container, to give views of the mice from all 4 sides. Mice were trained for 2 hours daily from Day 1 to Day 10 and were placed into the container 30 min before video recording to let the mice get used to the surroundings. For each 2-hour video, the itch-like scratching with the hindlimb and pain-like wiping with the forelimb was counted by an investigator who was blinded to the experimental design (13, 14).

### 2.4 Measurement of skin-fold thickness and PASI score

The image of each mice’s cheek skin was taken under brief anesthesia after the video recording in a standardized setup. The severity of the inflammation was rated for the amount of erythema, thickening and scaling, each assessed independently on the following scale: 0, none; 1, slight; 2, moderate; 3, marked; and 4, very marked. The PASI score was calculated as the sum of the erythema, scaling and thickening scores. The cheek fold thickness of each mouse was measured 3 times using a micrometre (Mitutoyo, Tokyo, Japan), and the mean was calculated.

### 2.5 Real-time quantitative polymerase chain reaction (RT-qPCR)

After the measure of cheek fold thickness, mice were sacrificed and the trigeminal ganglion (TG) from the treated side was immediately isolated. The mRNA was extracted using TRIzol reagents (ThermoFisher Scientific, Waltham, MA, USA) cDNA was synthesized from 100 µg of total RNA by Prime Script^TM^ RT reagent kit with gDNA Eraser (Takara, Tokyo, Japan). Each cDNA sample was amplified for the gene of interest and β-actin in a 15 µL reaction volume TB Green^TM^ Premix Ex Taq™ II (Takara, Tokyo, Japan). The PCR conditions were 95 °C for 30 s followed by 40 cycles of 95 °C for 5 s and 60 °C for 60 s. The whole procedure was performed according to the manufacturer’s instructions. The mRNA levels of all genes were normalized to the measured mRNA values of the housekeeping gene β-actin. The primers used are listed in Table 1.

**Table 1.**
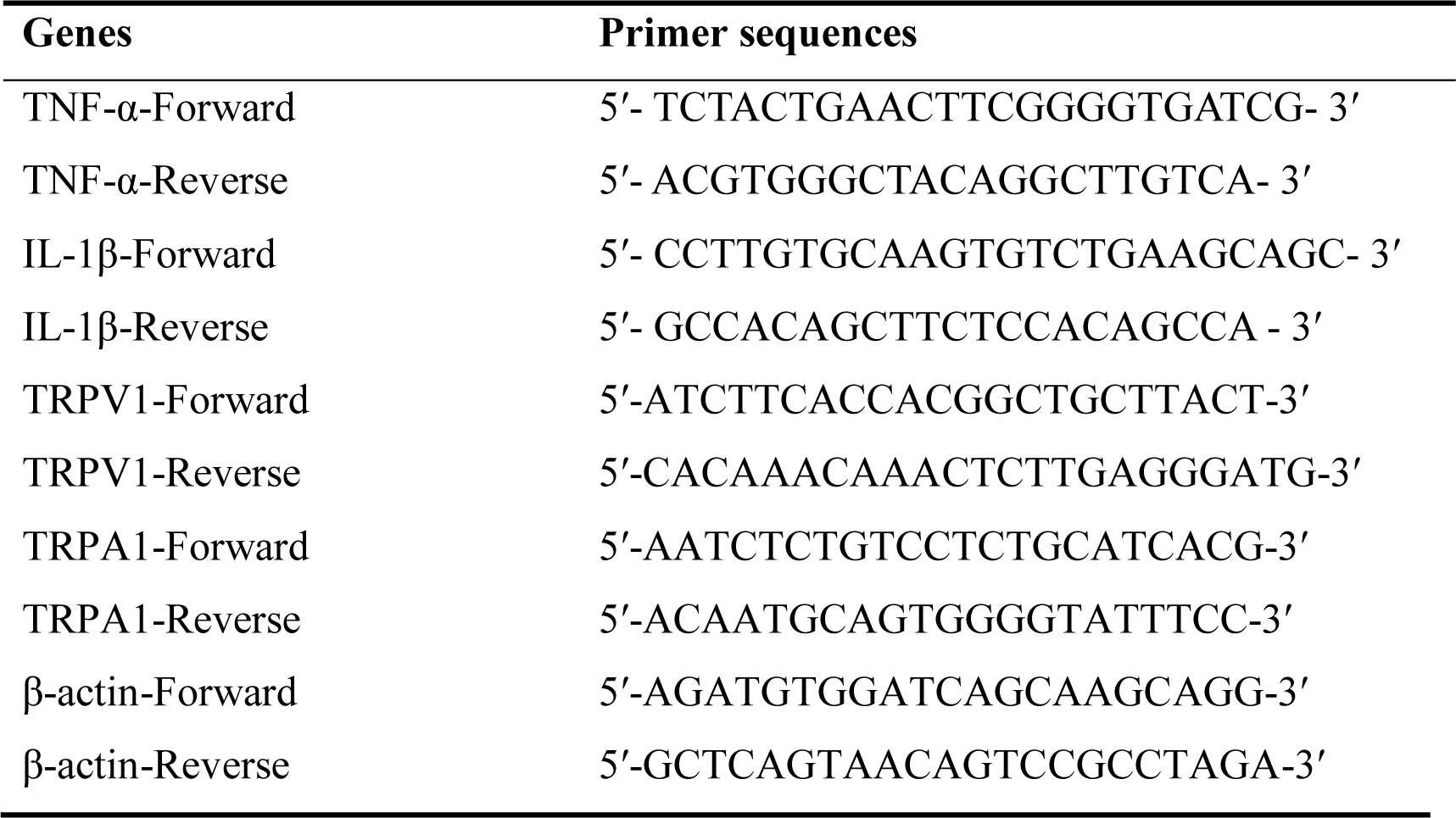
Primer sequences.

### 2.6 Western blotting

After the measure of cheek fold thickness, mice were sacrificed, and the cheek skin and TG from the treated side were immediately isolated. The total protein was extracted using Cell lysis buffer for Western and IP (Beyotime Biotechnology, Shanghai, China). 30 µg of each sample was added into 15% SDS gel. Protein separation was performed at room temperature with a voltage of 90 V for 30 min followed by 120V for 45-60 min, and protein transferring to the PVDF membrane was performed on ice with a current of 250 mA for 1.5 h. Afterwards, the PVDF membrane was washed 6 times with TBST on a shaking table at room temperature for 5 min each. Blocking was performed at room temperature using 5% skim milk powder in TBST on a shaking table for 1 h. Afterwards, the PVDF membrane was washed again 6 times with TBST on the shaking table at room temperature for 5 min each. The primary antibodies (rabbit polyclonal anti-TRPV1 1:1000 (NBP1-97417, Novus Biologicals, Littleton, CO, USA), rabbit polyclonal anti-TRPA1 1:2000 (NB100-98841, Novus Biologicals, Littleton, CO, USA) and mouse monoclonal β-actin 1:1000 (8H10D10, CST, Danvers, MA)) were incubated with PVDF respectively overnight at 4 ℃. After 6 times washing with TBST for 5 min each, HRP-conjugated secondary antibodies (included goat anti-rabbit 1:5000 (A0208, Beyotime Biotechnology, Shanghai, China) and goat anti-mouse 1:2000 (A0216, Beyotime Biotechnology, Shanghai, China) was incubated with PVDF membrane on the shaking table at room temperature for 2 h. After incubation, the PVDF membrane was washed with TBST 6 times, 5 min each. The detection was performed with the Super ECL detection reagent (Applygen, Beijing, China).

### 2.7 CAP and AITC-induced behavioral testing

50 C57BL/6 mice were randomly allocated into 5 groups: Control group, AITC group, AITC+SC group, CAP group and CAP+SC group, in CAP and AITC-induced behavior testing. The 10 mg/kg SC or the same volume of saline was administrated i.g. for the CAP+SC and AITC+SC group or other groups at 1 h before CAP and AITC, respectively. After adaptation for 30 min, 1 mg/kg CAP (Jiuding Chemical, Shanghai, China, dissolved in 7% Tween-80), 1% AITC (w/v, Chengdu Herbpurify Co., Ltd, Chengdu, China, dissolved in 7% Tween-80) and 10 µL 7% Tween-80 as the vehicle was intradermally administrated into mice cheek of CAP groups, AITC groups and control group respectively. In this assay, the video was recorded for 30 min following CAP and AITC injection. The pain-like behavior and itch-like behavior were measured as described in 2.3.

### 2.8 Molecular Docking

The AutoDock Vina in Chimera and ChimeraX were used for molecular docking simulations and visualization of SC, CAP and AITC into active binding sites of TRPA1 and TRPV1. The molecular structure of SC (ZINC3881784), CAP (ZINC1530575) and AITC (ZINC1687017) was adopted from ZINC (https://zinc.docking.org/).(15) The protein structure of TRPV1 (PDB ID: 3J5P) and TRPA1 (PDB ID: 3J9P) was adopted from RCSB Protein Data Bank (https://www.rcsb.org/).(16) Autodock Vina 1.1.2 in UCSF Chimera 1.17.3 was used for molecules and proteins preprocessing, docking box determination and docking simulations, according to a similar tutorial on https://vina.scripps.edu/. (17, 18) The docking results were visualized and atomic distances of hydrogen bonds were measured by ChimeraX 1.6.1. (19)

### 2.9 Statistical analysis

All the data was shown as mean±S.E.M. All statistical analyses were performed using GraphPad Prism 9 (GraphPad Software, La Jolla, CA, USA). The comparisons of quantitative data among groups were performed using one-way ANOVA, followed by Bonferroni corrections for testing between individual means. A value of *p* < 0.05 was considered statistically significant. All figures were processed in GraphPad Prism 9.

## 3 Results

### 3.1 SC reduced scratching and wiping behaviors in SADBE-induced ACD mice

Mice in the ACD group exhibited a significantly greater number of scratching bouts and wipes than in the control group (Figure 2). Compared with the ACD group, mice in the DEM group, CPZ group, and all SC groups exhibited fewer scratching bouts, while only SC groups exhibited fewer wipes (Figure 2). SC reduced mice wiping behavior in a dose-dependent manner, whereas 60 mg/kg and 30 mg/kg SC showed the same potency against scratching behavior, having however stronger effect than 10 mg/kg dose (Figure 2).

**Figure 2.**
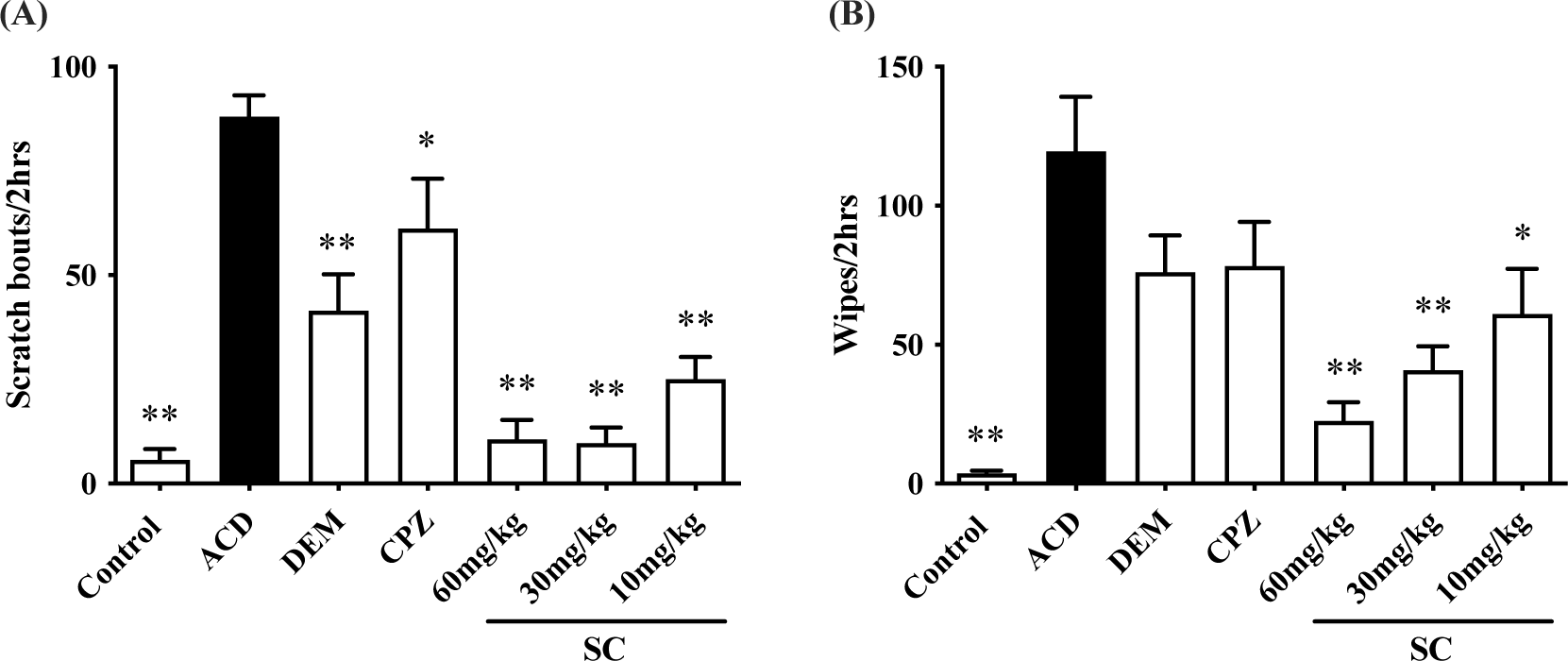
Effect of SC on relieving wiping and scratching on ACD mouse model. The mean of wipes (A) and the mean of scratching bouts (B) were evaluated for 2 h starting at 1 h after saline (control and ACD group), 5 mg/kg DEM (DEM group), 30 mg/kg CPZ (CPZ group), 60 mg/kg SC (60 mg/kg SC group), 30 mg/kg SC (30 mg/kg SC group), 10 mg/kg SC (10 mg/kg SC group) administration on ACD mice. \**p* < 0.05, \*\**p* < 0.01, compared with ACD group, error bars: S.E.M. n =10 mice /group.

### 3.2 SC improved the cheek skin inflammation in ACD mice

In the control group, we found no visual indications of redness, scaling or swelling of the cheek skin (Figure 3B). The ACD group demonstrated noticeable scaling, swelling and redness (Figure 3B). The skinfold thickness of the DEM group, 30 mg/kg SC group and 10 mg/kg SC group was lower than that of the ACD group, while there was no difference between the CPZ group, 60 mg/kg SC group and the ACD group (Figure 3A). Further analysis showed that the DEM group and SC groups had significantly lower PASI scores compared to the ACD group (Figure 3F). In the detailed scoring of the PASI scale, there was no difference between the DEM group, all SC groups and the ACD group in erythema scoring; the DEM group and 60 mg/kg SC group scored lower in scaling score; and all groups scored lower in thickening score compared with those of the ACD group (Figure 3C, 3D, 3E).

**Figure 3.**
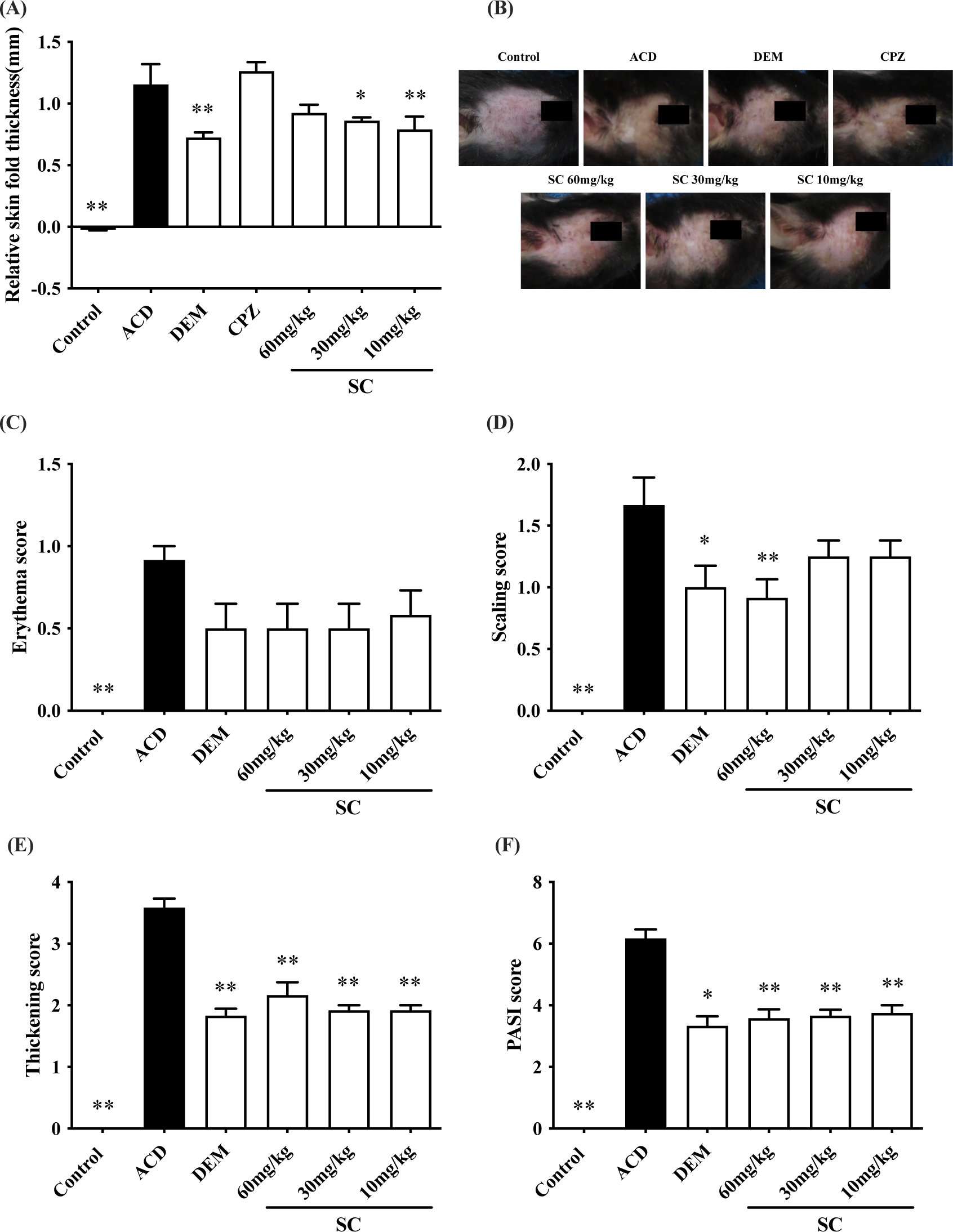
Effects of SC on thickness and severity of inflammation of the skin on ACD mouse model. The mean skin fold thickness (A) was measured after recording. (B) Exemplary photographs of the mouse cheek skin in each experimental group show the visible change in the different treatment groups. The erythema (C), scaling (D), and thickening (E) were separately scored on a scale from 0 (none) to 4 (very marked). The PASI score (F) was taken as the sum of the erythema, scaling and thickening scores. \**p* < 0.05, \*\**p* < 0.01 for significance compared with the ACD group, error bars: S.E.M. n=10 mice/group.

### 3.3 SC reduced inflammation by downregulating the expression of the TNF-α and IL-1β in the skin

Expression of TNF-α and IL-1β mRNA in mice cheek skin was significantly greater in the ACD group, compared with that of the control group (Figure 4A, 4B). Compared with the ACD group, DEM and SC significantly downregulated expression of TNF-α and IL-1β mRNA (Figure 4A, 4B). Western blotting revealed a significantly higher TNF-α and TRPV1 protein expression in the ACD group compared with the control group (Figure 4C, 4D). Compared with the ACD group, DEM and SC decreased TNF-α and IL-1β protein expression (Figure 4C, 4D).

**Figure 4.**
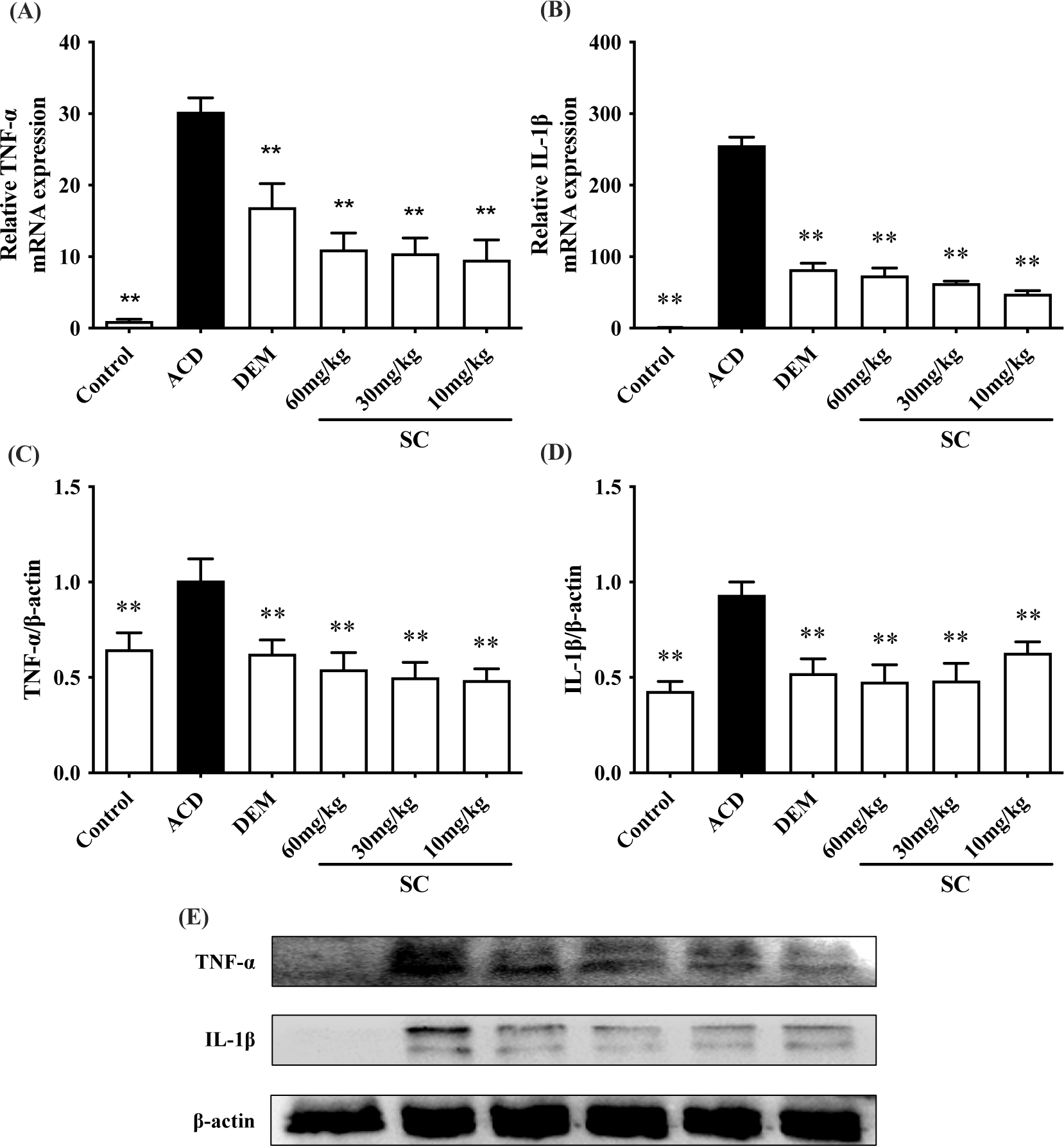
Effects of SC on proinflammatory factors in the skin on ACD mouse model. The mRNA expression of TNF-α (A) and IL-1β (B) were obtained from cheek skin for different groups of mice. The mRNA data are presented as 2^-ΔΔCt^ values relative to β-actin. Protein expression of TNF-α (C) and IL-1β (D) and representative of western blots (E) respectively. \**p* < 0.05, \*\**p* < 0.01 for significance compared with the ACD group, error bars: S.E.M. n = 3-6 mice/group.

### 3.4 SC downregulated the expression of the TRPA1 & TRPV1 in TG

Expression of TRPA1 and TRPV1 mRNA in TG was significantly greater in the

ACD group, compared with that of the control group (Figure 5A, 5B). Compared with the ACD group, CPZ and SC significantly downregulated TRPV1 and TRPA1 mRNA expression. (Figure 5A, 5B). At the protein level, significantly higher TRPA1 and TRPV1 protein expression was identified in the ACD group compared with the control group (Figure 5C, 5D). Compared with the ACD group, CPZ had no effect, whereas SC decreased TRPA1 and TRPV1 protein expression (Figure 5C, 5D).

**Figure 5.**
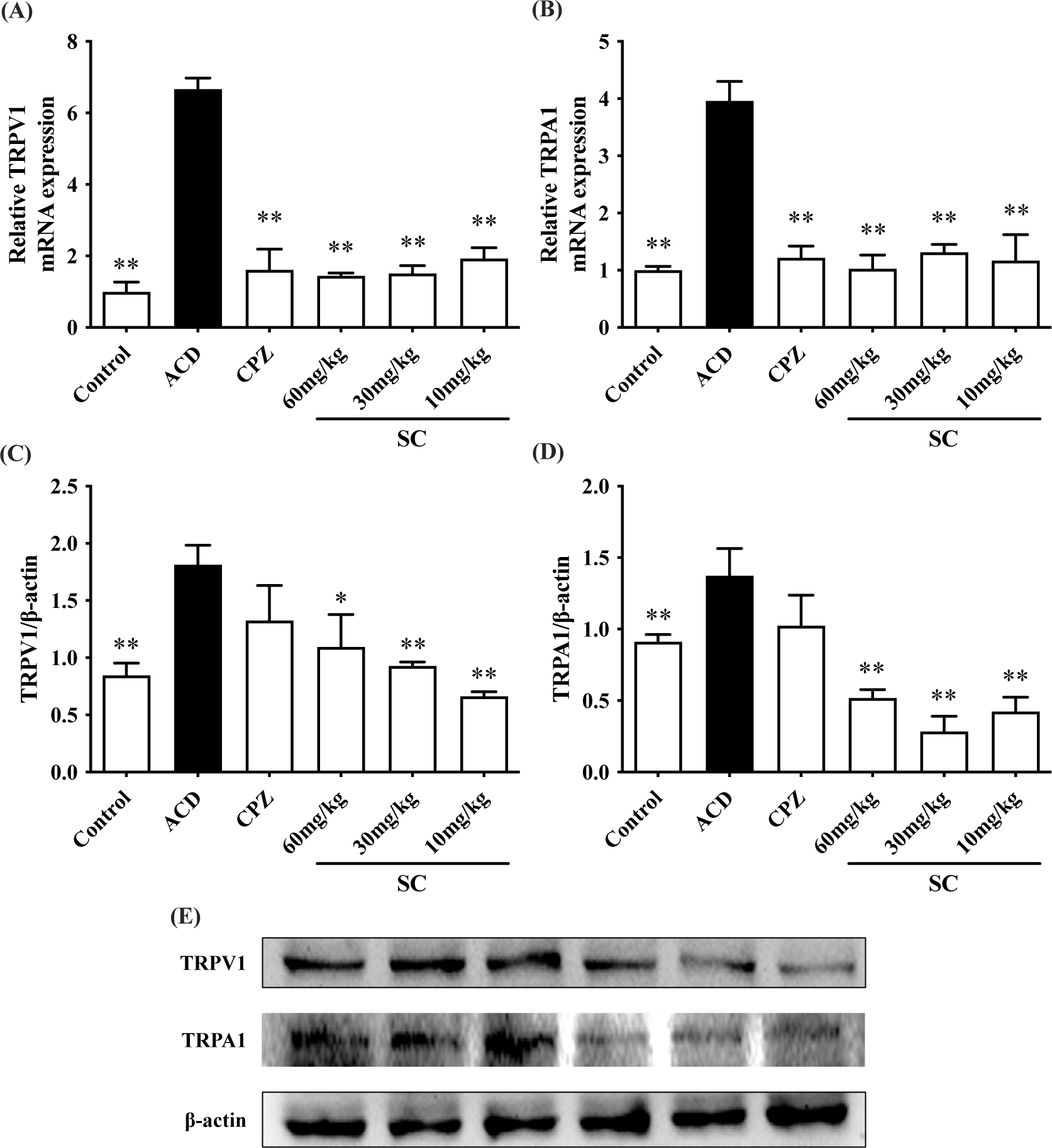
Effects of SC on TRP channels expression in TG of ACD mouse model. The mRNA expression levels of TRPV1 (A) and TRPA1 (B) were obtained from TG for different groups of mice. The mRNA data are presented as 2^-ΔΔCt^ values relative to β-actin. The protein expression of TRPV1 (C) and TRPA1 (D), and exemplary images of western blotting (E). \**p* < 0.05, \*\**p* < 0.01 for significance compared with the ACD group, error bars: S.E.M. n = 3-5 mice/group.

### 3.5 SC reduced scratching and wiping behavior induced by AITC and CAP

To determine whether SC targets TRPA1 and/or TRPV1, 10 mg/kg SC was administrated i.g. before treatment with AITC or CAP. The numbers of scratching bouts and wipes were significantly increased after the AITC administration (Figure 6A, 6B), while only wipes were significantly increased after CAP administration (Figure 6E, 6F). The wipes peaked at 5 min after ATIC and CAP administration, while the scratching bouts peaked at 25 min after ATIC administration (Figure 6D, 6H, 6C). The scratching bout was not significantly induced by CAP (Figure 6G). SC significantly decreased scratching bouts induced by AITC and wipes induced by AITC and CAP (Figure 6A, 6B, 6F).

**Figure 6.**
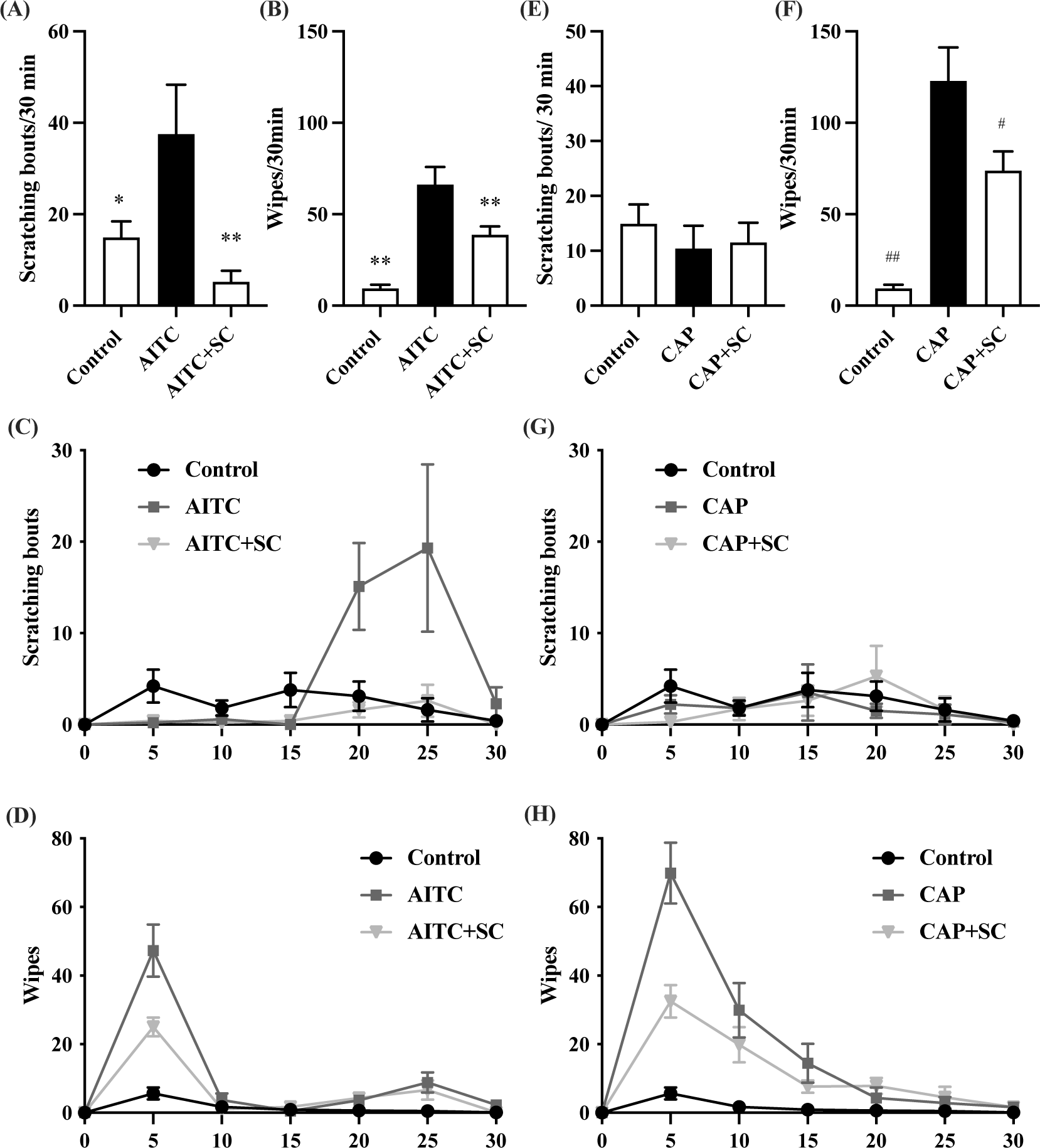
Effect of SC in AITC-induced and CAP-induced wiping and scratching behavior. The numbers of scratching bouts (A) and wipes (B) were evaluated for 30 min after AITC injection at 1 h after SC administration. The numbers of scratching bouts (E) and wipes (F) were evaluated for 30 min after CAP injection at 1 h after SC administration. The time course of scratching bouts (C) and wipes (D) after AITC injection at 1 h after administration. The time course of scratching bouts (G) and wipes (H) after AITC injection at 1 h after administration. \**p* < 0.05, \*\**p* < 0.01 for significance compared with AITC group, ^#^*p* < 0.05, ^##^*p* < 0.01 for significance compared with CAP group, error bars: S.E.M, n=10 mice/group.

### 3.6 SC as a competitive inhibitor of TRPA1 and TRPV1

To further investigate the relationship between SC, AITC, CAP and TRPV1, TRPA1, molecular docking simulation was performed *in silico* by Autodock Vina and Chimera/ChimeraX, with small molecules, or ligands, of SC, AITC and CAP, and target protein, or receptors, of TRPV1 and TRPA1 tetramers. The highest binding energies estimated between SC and TRPV1 or TRPA1 were higher than that of AITC, but lower than that of CAP. All ligands can form hydrogen bonds with receptors except for AITC and TRPV1 (Table 2, Figure 7). The atom distance between ligands and receptors estimated between SC and TRPV1 or TRPA1 was lower than that of AITC but higher than that of CAP.

**Figure 7.**
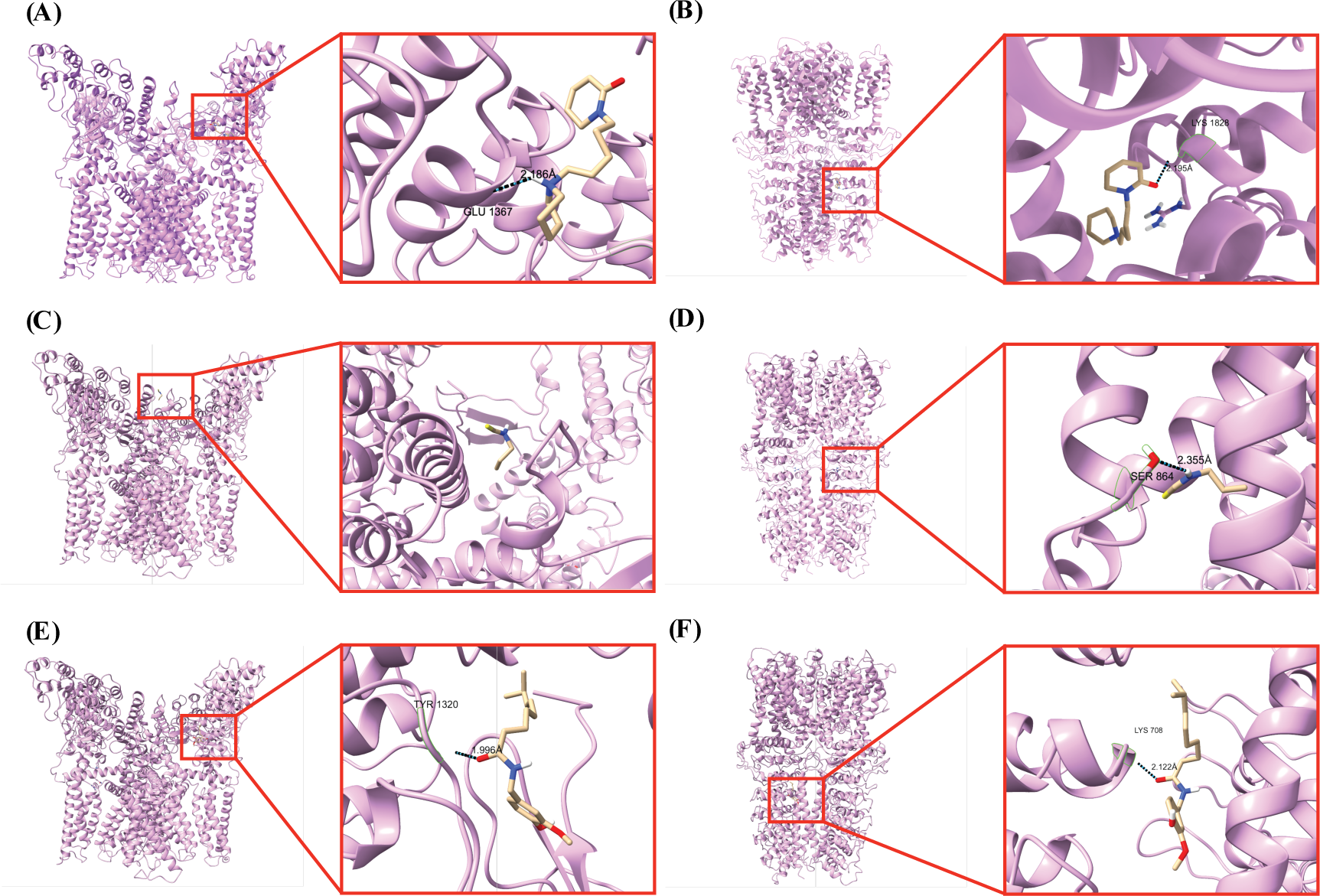
Estimated optimal binding conformations by docking simulation of ligands (small molecules, in yellow) to receptors (proteins, in purple). SC was docked onto (A) TRPV1 and (B) TRPA1; AITC was docked onto (C) TRPV1 and (D) TRPA1; CAP was docked onto (E) TRPV1 and (F) TRPA1. The distance of the hydrogen bonds between ligands and receptors was calculated.

**Table 2.** Estimated binding energy by molecular docking.

| Ligand | Receptor | KCal/mol | Binding Type | Binding Site | Atoms Distance(Å) |
| --- | --- | --- | --- | --- | --- |
| SC | TRPA1 | -6.1 | Hydrogen | Ser364 | 2.186 |
|  | TRPV1 | -6.4 | Hydrogen | Glu1367 | 2.195 |
| AITC | TRPA1 | -3.5 | Hydrogen | Ser864 | 2.355 |
|  | TRPV1 | -3.4 | Van der Waals |  |  |
| CAP | TRPA1 | -6.9 | Hydrogen | Lys708 | 1.996 |
|  | TRPV1 | -7.1 | Hydrogen | Tyr1320 | 2.122 |

To further investigate the relationship between SC, AITC, and CAP when binding to TRPV1 and TRPA1, the competition of docking results was visualized by ChimeraX. SC may compete with AITC by binding to TRPV1 monomer at SER284, GLU287 in the Linker domain, LEU438, ARG 442 in the S4-S5 linker, and GLN562 in the TRP domain (Figure 8A), and by binding to TRPA1 monomer at sites of ASN145, LYS146, ARG147, LYS148 in AR16 and LYS528 in the C-terminal coiled-coil (Figure 8C), or disturbing AITC binding to ILE234, GLU235, ARG238 in the Pre-S1 helix and LEU479, SER521 in the TRP-like domain by Van der Waals. SC may compete with CAP by binding to TRPV1 monomer at GLU287, ARG299 in the Linker domain, SER397 in the S3, ARG442 in the S4-S5 linker, and LYS576 in the TRP domain (Figure 8B).

**Figure 8.**
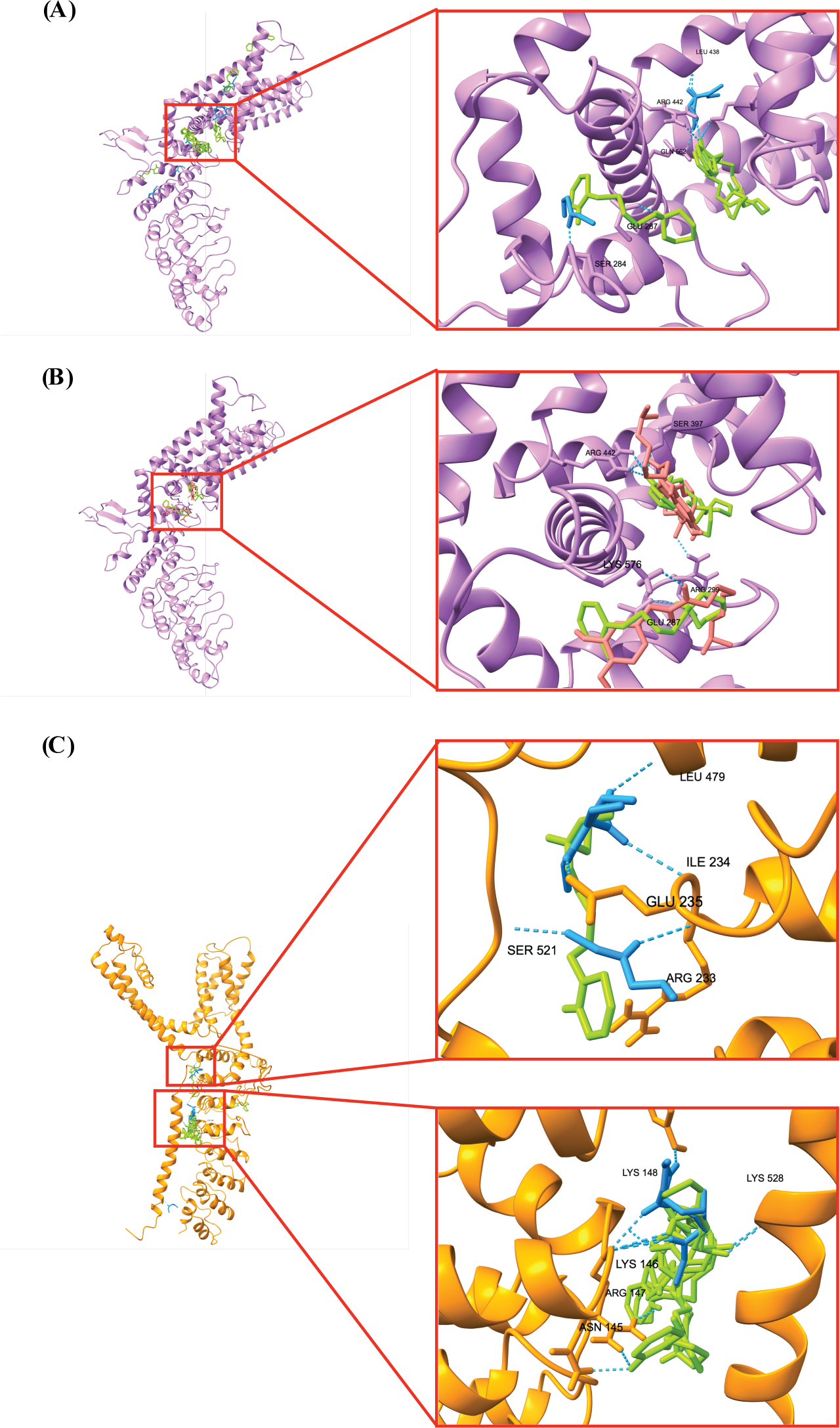
The potential competition between SC and AITC/CAP on TRPV1 and TRPA1 monomers. (A) SC (green) and AITC (blue), and (B) SC and CAP (red) competed in one protein pocket of TRPV1 (purple); (C) SC (green) and AITC (blue) competed on different binding sites in two protein pockets of TRPA1 (orange).

## 4. Discussion

Spontaneous itch and pain are of paramount clinical relevance being the main complaint of chronic itch and pain patients (3). SC was screened from SFR by our research group which showed a binding ability to DRG cells (11). In the present study, the anti-pruritic and analgesic effects of SC were tested in a SADBE-induced ACD mouse model manifesting both itch and pain. Our results revealed that SC possesses strong anti-pruritic and analgesic effects. Expression of mRNA and protein of TRPA1 and TRPV1 in SC treatment groups was decreased compared to the ACD group. Furthermore, *in vivo* experiments showed that SC reduces AITC-induced wiping and scratching behavior and CAP-induced wiping behavior thus indicating that SC inhibits both TRPV1 and TRPA1. Molecular docking revealed the potential of SC as a competitive inhibitor of both TRPV1 and TRPA1, compared with AITC and CAP. Therefore, we posit that SC can relieve itch and pain in ACD by downregulating expression and inhibiting the function of TRPV1 and TRPA1.

The patterns of peripheral and central sensitization linked to chronic itch and pain are remarkably similar (3). In the clinic, ACD-associated itch symptoms are often presented with repeated long-lasting episodes and are sometimes accompanied by pain sensation (20). SADBE is most commonly used for the treatment of pelade and warts but often causes ACD with severe itch after frequent applications, and hence SADBE is commonly used in ACD studies (21). Consistent with clinical observations, SADBE effectively induces ACD in mice, which display significant spontaneous scratching and wiping behavior (22). In the present study, SADBE was used to induce the ACD mouse model; SABDE increased scratching bouts and wipes significantly as has been described previously (23).

SFR is widely used in relieving inflammatory itch and pain. Oxymatrine, screened from SFR together with SC by our research group, has already been tested and demonstrated anti-pruritic and anti-inflammatory effects in ACD mice (11, 12). It was previously shown that SC possessed significant anti-nociceptive activity in thermally and chemically induced mouse pain models, and anti-inflammatory activity on carrageenan-induced rat hind paw edema, xylene-induced mouse ear edema and acetic acid-induced mouse vascular permeation (24). In the present study, the anti-pruritic effect of SC was additionally evaluated and all these three effects were tested on the same mouse model, with 10 mice for each group to reduce the individual difference. Our results revealed that SC significantly reduces scratching bouts and wipes in the ACD mice, thus showing a strong anti-pruritic and analgesic effect. The PASI score and skin fold thickness measurement showed that SC also relieved the SADBE-induced inflammation reaction. TNF-α is the major cytokine involved in chronic inflammation inducing IL-1β and other interleukins for inflammatory signaling cascades. In the present study, SC downregulated mRNA and protein expression of TNF-α and IL-1β as good as dexamethasone in the ACD model.

TRPA1 and TRPV1 are both involved in inflammatory pain and were suggested to mediate histamine-independent and histamine-dependent itch, respectively (7). Both TRPA1 and TRPV1 are required for generating spontaneous scratching in the SADBE-induced ACD mouse model by directly promoting the excitability of pruriceptors (25). In the present study, we found that TRPA1 and TRPV1 in TG were upregulated and increased after the SADBE application, consistent with previous data from the dorsal root ganglia.

TRPA1^−/−^ mice manifest fewer pain responses when injected with bradykinin or ATIC (26). Mice lacking TRPA1 display reduced scratching and skin lesion severity, highlighting a role for TRPA1 in acute and chronic itch, and suggesting that TRPA1 antagonists may be useful for treating various pathological itch conditions (27). Nevertheless, mice lacking TRPA1 and subjected to histamine-independent itch modeling showed less reduction of scratching response to cutaneous application of borneol (28). Expression of TRPA1 was upregulated in patients who suffer from severe itch (29). The mRNA of TRPA1 was upregulated in neuropathic pain whereas the protein of TRPA1 was increased in inflammatory pain in dorsal root ganglia (30, 31). TRPA1 also contributed to pruritus in allergic contact dermatitis (32, 33). Our results demonstrated that SC downregulates the TRPA1 expression and reduces the scratching and wiping behaviors induced by ATIC, similar to HC-030031 (34). Molecular docking showed lower binding energy and a smaller atomic distance of hydrogen bond between SC and TRPA1, indicating that SC could be a potential competitive inhibitor of TRPA1, compared with AITC. Thus, we suggest that SC efficiently inhibits the function and downregulates the expression of TRPA1 to relieve itch and pain in diseases such as ACD.

TRPV1 is expressed at the highest level in peripheral sensory neurons involved in the perception of pain (35). The first evidence of the role of TRPV1 in peripheral pain processing arose from observations made in mice injected with the TRPV1 antagonist CPZ (36, 37). TRPV1 contributes to both acute and chronic itch conditions (38). CPZ pretreatment reduced the scratching bouts induced by the subcutaneous injection of histamine into mice’s necks (39). The anti-pruritic and analgesic effect of CPZ was also observed in our present study in the SADBE-induced ACD mouse model. A point mutation in the coding of the S4-S5 linker in the *Trpv1* gene reduced TRPV1 phosphorylation and cell membrane recruitment, successively reducing pain and histamine-dependent itch (40). Oleic acid, a natural compound, also inhibits TRPV1 thus modulating itch and pain (41). We also showed that SC downregulated the mRNA and protein expression of TRPV1. Our behavioral tests showed that CAP-induced wiping was reduced by pretreatment of SC, hinting that SC could inhibit TRPV1. Molecular docking showed competitive binding energy and atomic distance of hydrogen bond between SC and TRPV1, indicating that SC may be a potential competitive inhibitor of TRPV1 Thus, we suggest that SC could efficiently inhibit and downregulate the expression of TRPV1 to relieve itch and pain manifested in diseases such as ACD.

Molecular docking simulations demonstrated that SC is competing on the intracellular AR16, the C-terminal coiled-coil, the membrane Pre-S1 helix and the TRP-like domain in the TRPA1 monomer with AITC, and competing on the Linker domain, the S4-S5 linker, and the TRP domain in the TRPV1 monomer with AITC and CAP. Although the amino acid numbers differ, the binding pocket around the S4-S5 linker in TRPV1 of SC competing with AITC or CAP simulated in this study is quite similar to the previously published simulation of CAP binding to TRPV1, referring to the TRPV1 structure identified (42, 43). While the major simulations of AITC and SC demonstrated lower binding energy to the intracellular AR16 or the Linker domain in the N-terminus, the disturbance of SC in the Pre-S1 helix and the TRP-like domain seems more convincing, compared to the simulation of DNCB and DNFB, agonists of TRPA1 to trigger itch as AITC, binding to TRPA1 in protein pockets around the Linker domain and the TRP-like domain (44). Further studies focusing on protein pockets identified to testify whether SC is a competitive inhibitor of TRPV1 and TRPA1 are needed.

In conclusion, SC demonstrated a strong anti-inflammatory, anti-pruritic and analgesic effect by targeting TRP channels in TG. These findings suggest that SC might have significant therapeutic potential in treating inflammatory itch and pain.

## 5. Conflict of interest

The authors declare no conflicts of interest.

## 6. Author contributions

H.Z. and Z.Z. collected and analyzed the data. H.Z., P. L. collected the samples. The mice cheeks were scored by D.Z. and Z.Z. The video analysis was done with assistance from P.L. and D.Z. H.N. and A.V. designed the experiments. The paper was written by H.Z., and modified by H.N and A.V. The research is supervised by H.N.

## 7. Acknowledgements

This work was supported by grants from the Chinese National Natural Science Foundation to H.N. (No. 81373993 & No. 81861138042).

## Notes

### Competing Interest Statement

The authors have declared no competing interest.

### Summary of Updates

Alexei Verkhratsky as the corresponding author updated. Table 1 updated with missing primers.

